# SARS-CoV-2 infection alkalinizes the ERGIC and lysosomes through the viroporin activity of the viral envelope protein

**DOI:** 10.1101/2022.10.04.510830

**Authors:** Wen-An Wang, Amado Carreras Sureda, Nicolas Demaurex

## Abstract

The coronavirus SARS-CoV-2, the agent of the deadly COVID-19 pandemic, is an enveloped virus propagating within the endocytic and secretory organelles of host mammalian cells. Enveloped viruses modify the ionic homeostasis of organelles to render their intra-luminal milieu permissive for viral entry, replication, and egress. Here, we show that infection of Vero E6 cells with the delta variant of the SARS-CoV-2 alkalinizes the endoplasmic reticulum-Golgi intermediate compartment (ERGIC) as well as lysosomes, mimicking the effect of inhibitors of vacuolar proton ATPases. We further show the envelope protein of SARS-CoV-2 accumulates in the ERGIC when expressed in mammalian cells and selectively dissipates the ERGIC pH. This viroporin effect is not associated with acute cellular toxicity but is prevented by mutations within the channel pore of E. We conclude that the envelope protein acts as a proton channel in the ERGIC to mitigate the acidity of this intermediate compartment. The altered pH homeostasis of the ERGIC likely contributes to the virus fitness and pathogenicity, making the E channel an attractive drug target for the treatment of COVID-19.

## Introduction

The severe acute respiratory syndrome-coronavirus 2 (SARS-CoV-2), the agent of the dramatic ongoing COVID-19 pandemic, is the third coronavirus to cause severe respiratory disease in humans. SARS-CoV-2 propagates more efficiently than the highly pathogenic human coronaviruses SARS-CoV and MERS-CoV that emerged in 2002 and 2012, respectively (Gates, 2020). Up to 20% of hospitalized infected post-COVID patients require respiratory support for acute respiratory distress syndrome (ARDS) triggered by a cytokine storm. Lung damage, intravascular coagulation, and severe T cell lymphopenia characterize late-stage SARS-CoV-2 infection, with viral toxicity and hyper-inflammation both contributing to pathogenicity. A significant proportion of infected patients suffer from multisystemic effects cumulating into long-COVID symptoms that can be highly debilitating (Silva Andrade et al., 2021). Current therapeutic strategies combine antiviral drugs and immune modulators (Harrison, 2020) but are hampered by a lack of information on the molecular and cellular mechanisms of SARS-CoV-2 infection, putting intense pressure on prevention measures that rely on the worldwide distribution and administration of safe and effective vaccines.

The SARS-CoV-2 genome encodes for 4 structural proteins required to produce a complete infectious viral particle and 16 non-structural proteins that drive viral replication and contribute to viral pathogenicity. The spike (S) structural protein mediates virus entry via attachment to the angiotensin-converting enzyme (ACE2) receptor after priming by cellular proteases (Hoffmann et al., 2020). The nucleocapsid (N) protein binds the single-stranded, positive-sense viral RNA genome and organizes its replication. The membrane (M) and envelope (E) proteins drive virus assembly and budding. Once bound to cell surface ACE2 receptors the S protein requires proteolytic cleavage by host cell proteases to drive the fusion of viral and cellular membranes. Depending on protease availability and cell type, the S protein can be cleaved at the plasma membrane (PM) by the serine protease TMPRSS2 (Hoffmann et al., 2020) or in endosomes by the cysteine proteases cathepsins B and L (Mingo et al., 2015; Smieszek et al., 2020). Direct PM entry is more efficient and is the preferred route in lung cells expressing TMPRSS2 (Hoffmann et al., 2020). Endocytosis is the default entry route and requires an acidic pH in the lumen of endosomes to activate cathepsin L (Simmons et al., 2005). Membrane fusion releases the viral genome into the host cell cytoplasm, generating viral proteins required for RNA synthesis and for the formation of intracellular double-membrane structures derived from the endoplasmic reticulum (ER) that serve as scaffold for viral replication and as protection from antiviral host cell responses (Knoops et al., 2008).

Upon translation, the S, E, and M structural proteins insert into the membrane of the ER and drive viral assembly in the ER-Golgi intermediate compartment (ERGIC). The virions then bud into the ERGIC lumen and accumulate in large vesicles to reach the PM and egress (Ruch and Machamer, 2012; Ulasli et al., 2010). Assembled β-Coronaviruses were also shown to egress through lysosomal exocytosis (Ghosh et al., 2020). SARS-COV-2 E and M redirect S to the ERGIC and are required for the optimal production of viral-like particles (Boson et al., 2021). The E and M proteins interact to drive membrane bending and scission (Ruch and Machamer, 2012), and exogenous expression of E generates tubular convoluted membranes mimicking those of infected cells (Raamsman et al., 2000). Combined E and M expression is sufficient to generate viral-like particles (Baudoux et al., 1998) and deleting the E gene reduces SARS-CoV infectivity, leading to intracellular accumulation of virions with aberrant material (DeDiego et al., 2007). Most of the E protein, however, does not incorporate in the virus, suggesting that it sustains infection by altering host cell functions.

The SARS-CoV-2 E protein (UniProtKB:P59637) is a small, 75 amino acid protein, containing a predicted amphipathic transmembrane alpha-helix followed by a cluster of positively charged residues, two signature motifs characteristic of viral ion channels, or viroporins (Hyser and Estes, 2015). The E protein is the most conserved of SARS-CoV structural proteins, with 100% identity between SARS-CoV-2 and a CoV isolated from a Malayan pangolin, the likely intermediate host in the COVID-19 pandemic (Xiao et al., 2020). All CoV E proteins studied so far have ion channel activity and are inhibited by micromolar concentrations of hexamethylene amiloride (HMA) (Surya et al., 2015; Verdia-Baguena et al., 2012; Wilson et al., 2006). NMR spectroscopy of SARS-CoV E revealed a pentameric channel (Pervushin et al., 2009; Surya et al., 2018) (PDB ID: 5X29) with residues Asp15 and Val25 contributing to oligomerization and ion conductance (Pervushin et al., 2009). The E protein of avian infectious bronchitis increases the pH of the Golgi (Westerbeck and Machamer, 2019) and the E protein of SARS-CoV functions as a Ca^2+^ permeable channel in artificial membranes (Nieto-Torres et al., 2015). The Ca^2+^ channel activity of SARS-CoV promoted viral replication and pathogenesis by disrupting the host Ca^2+^ signalling pathways (Hyser and Estes, 2015), and loss of channel function reduced virus fitness and pathogenicity (Regla-Nava et al., 2015). The structure of the transmembrane domain of the SARS-COV-2 E protein was determined by solid-state NMR spectroscopy at 2.1-Å resolution (Mandala et al., 2020). The transmembrane domain reconstituted into ERGIC-mimetic lipid bilayers formed a five-helix bundle surrounding a narrow and partially dehydrated pore, consistent with its predicted channel function. The viroporin function of the E protein of SARS-CoV-2 was recently established by electrophysiological recordings. Membrane currents carried by Na+ and K+ were recorded in planar lipid bilayers containing recombinant E (Xia et al., 2021) and in cells ectopically expressing E lacking its ER retention sequence and bearing a Golgi export sequence to ensure its expression at the plasma membrane (Cabrera-Garcia et al., 2021). This indicates that the E protein of SARS-CoV-2 forms an ion channel permeable to monovalent cations. The currents were sensitive to changes in extracellular pH (corresponding to changes in the pH of the ERGIC lumen) and expression of a tagged SARS-CoV-2 E increased the pH reported by an acidophilic dye (Cabrera-Garcia et al., 2021), indicating that E dissipates the pH of acidic organelles, as expected from its viroporin activity.

Here, we study the impact of SARS-CoV2 infection on the pH homeostasis of intracellular compartments of its host mammalian Vero E6 cells. By recording the pH within the cytoplasm, ERGIC and lysosome with genetically encoded pH probes after SARS-CoV2 infection or following ectopic expression of the SARS-CoV-2 E protein, we show that viral infection deacidifies both the ERGIC and lysosomes while expression of the E protein alone increases the ERGIC pH, an effect that was not observed with an E protein bearing pore mutations. The E protein therefore acts as a viroporin to alkalinize the ERGIC during viral infection, and likely contributes to virus pathogenicity.

## Results

### Validation of a genetically-encoded pH indicator targeted to the ERGIC

To assess whether viral infection with SARS-Cov2 alters the pH of the ERGIC lumen, we fused the ERGIC-transmembrane protein Sec22b to the ratiometic pH reporter probe pHluorin (Sec22b-rpHl) (Fig. 1A). When expressed in Vero E6 cells, Sec22b-rpHl decorated punctate perinuclear structures that colocalized extensively with ERGIC-53 immunoreactivity (Hauri et al., 2000), validating the proper targeting of the pH probe (Fig. 1A). Calibration on a high-resolution fluorescence microscope showed that the fluorescence ratio of Sec22b-rpHl increased 3.4-fold in Vero cells as pH was equilibrated from 5.5 to 8.0 with ionophores, with a pKa of 6.81 well resolved on a log-log pH titration fit (Fig. S1A). Nearly identical calibration curves were obtained in cells transfected with the cytosolic rpHl and the ERGIC Sec22b-rpHl, grown on 96-wells plates and imaged on an automated microscope placed in a biosafety level 3 (BSL3) laboratory (Fig. S1B). The calculated ERGIC pH of cells imaged within the BSL3 isolator was slightly more acidic than their cytosolic pH (pH_cyto_=7.27±0.32 vs. pH_ERGIC_=7.16±0.19, Fig. 1B). Inhibition of the vacuolar H^+^-ATPase with concanamycin A (ConA) increased ERGIC pH by 0.13 unit without affecting the cytosolic pH (Fig. 1C), indicating that the mildly acidic pH of the ERGIC reflects proton pumping by V-ATPases. These data validate sec22b-rpHl as a reliable quantitative reporter of the ERGIC luminal pH and reveals that this compartment is acidified by vacuolar proton ATPases.

**Figure 1.**
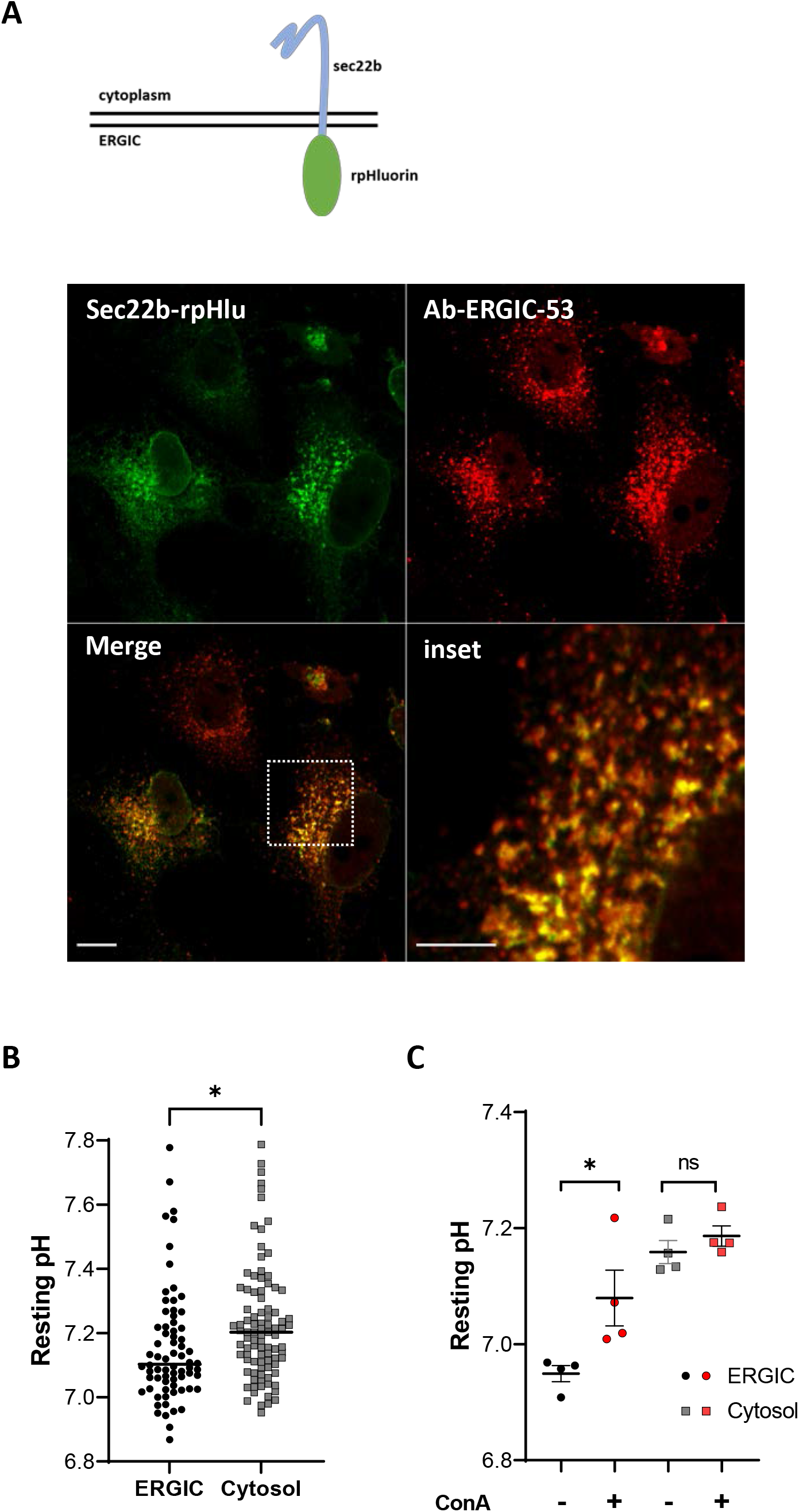
Recording the pH of the ERGIC lumen in Vero E6 cells. (A) Representative fluorescence images of VeroE6 cells transiently expressing Sec22b-rpHluorin (green) and stained with an anti-ERGIC53 antibody (red). Bars size 10 μm. Sketch shows the features of the Sec22b-rpHluorin construct used to measure ERGIC pH. (B) Calibrated pH values reported by the rpHluorin probe in the ERGIC and cytosol of VeroE6 cells. Each dot shows the average pH of 40-50 cells from 2 independent experiments, each with 30-50 image fields, lines show median values. *p<0.05, two-tailed unpaired Student’s t test. (C) Effect of Concanamycin A (ConA, 1 μM, 10 min) on the resting cytosolic and ERGIC pH of Vero E6 cells. Each dot shows the averaged pH of 15-50 cells from 4 replicates in one of two independent experiments, lines show mean±SD. *p<0.05, ns, non-significant, ordinary one-way ANOVA.

### SARS-Cov2 infection prevents the acidification of the ERGIC

Next, we measured pH_cyto_ and pH_ERGIC_ in Vero E6 cells infected with increasing MOIs of the delta SARS-CoV2 virus for 24 hours. Viral infection did not alter the expression levels and subcellular distribution of the Sec22b-rpHluorin probe (Fig. 2A). The pH_cyto_ was stable as the viral load increased, ranging from 7.15±0.03 at MOI 0 to 7.19±0.07 at MOI 1 (Fig. S2A) and remained insensitive to ConA (Fig. S2B). In contrast, pH_ERGIC_ was significantly higher at MOIs above 0.5, increasing from 6.96±0.03 at MOI 0 to 7.08±0.08 at MOI 1 (Fig. 2B), and the ERGIC alkalinization evoked by ConA was not observed in infected cells (Fig. 2C). These data indicate that infection with the SARS-Cov2 delta virus dissipates the acidic ERGIG pH maintained by V-ATPases.

**Figure 2.**
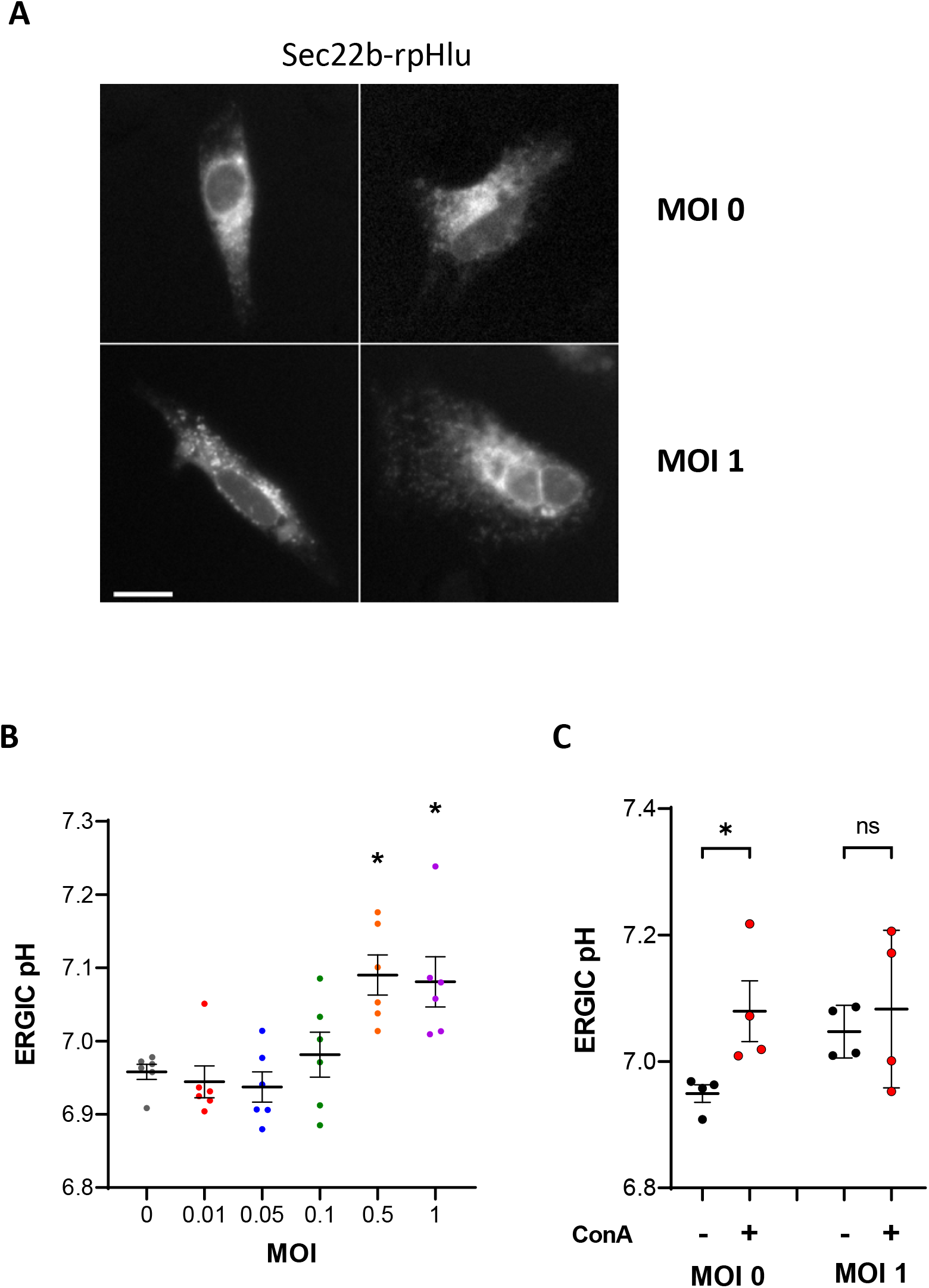
Effect of SARS-CoV-2 infection on the ERGIC pH of Vero E6 cells. (A) Representative fluorescence images (λex=488, λex=510 nm) of Vero E6 cells expressing Sec22b-rpHl and infected with 0 or 1 MOIs of the Delta SARS-CoV2 virus. Bar size 10 μm. (B) pH values reported by the ERGIC probe in Vero E6 cells infected with different MOIs of the Delta SARS-CoV2 virus. Each dot represents the average of 10-30 cells from 3 independent experiments performed in duplicates, lines show mean±SD. *p<0.05, ns, non-significant, ordinary one-way ANOVA. (C) ERGIC pH of Vero E6 cells infected with 0 (data from Fig. 1C) and 1 MOIs of the Delta SARS-CoV2 virus and treated or not with ConA (1μM, 10min). Each dot shows the averaged pH of 15-50 cells from 4 replicates in one of two independent experiments, lines show mean±SD. *p<0.05, ns, non-significant, ordinary one-way ANOVA.

### SARS-Cov2 infection mitigates lysosomal acidification

Assembled β-Coronaviruses exit through lysosomal exocytosis and SARS-Cov2 infection was reported to prevent lysosomal acidification (Cabrera-Garcia et al., 2021; Ghosh et al., 2020). We therefore measured the lysosomal pH (pHlyso) by exposing Vero E6 cells overnight to dextran particles labeled with Oregon Green (OGDx). Calibration indicated that OGDx fluorescence increased 4-fold in the pH range 4 to 6 (Fig. S3). Infection with increasing MOIs of the Delta SARS-CoV2 virus for 24 hours did not alter the OGDx loading pattern but increased OGDx fluorescence intensity (Fig. 3A), corresponding to an increase in pHlyso from 5.22±0.10 at MOI 0 to 5.83 ±0.10 at MOI1 (Fig. 3B). Moreover, ConA significantly increased pHlyso at MOI 0 but not at MOI 1 (Fig. 3C), consistent with a reduced contribution of V-ATPases in infected cells. These data indicate that infection with the delta SARS-Cov2 variant also mitigates the acidification of lysosomes by counteracting the activity of V-ATPases.

**Figure 3.**
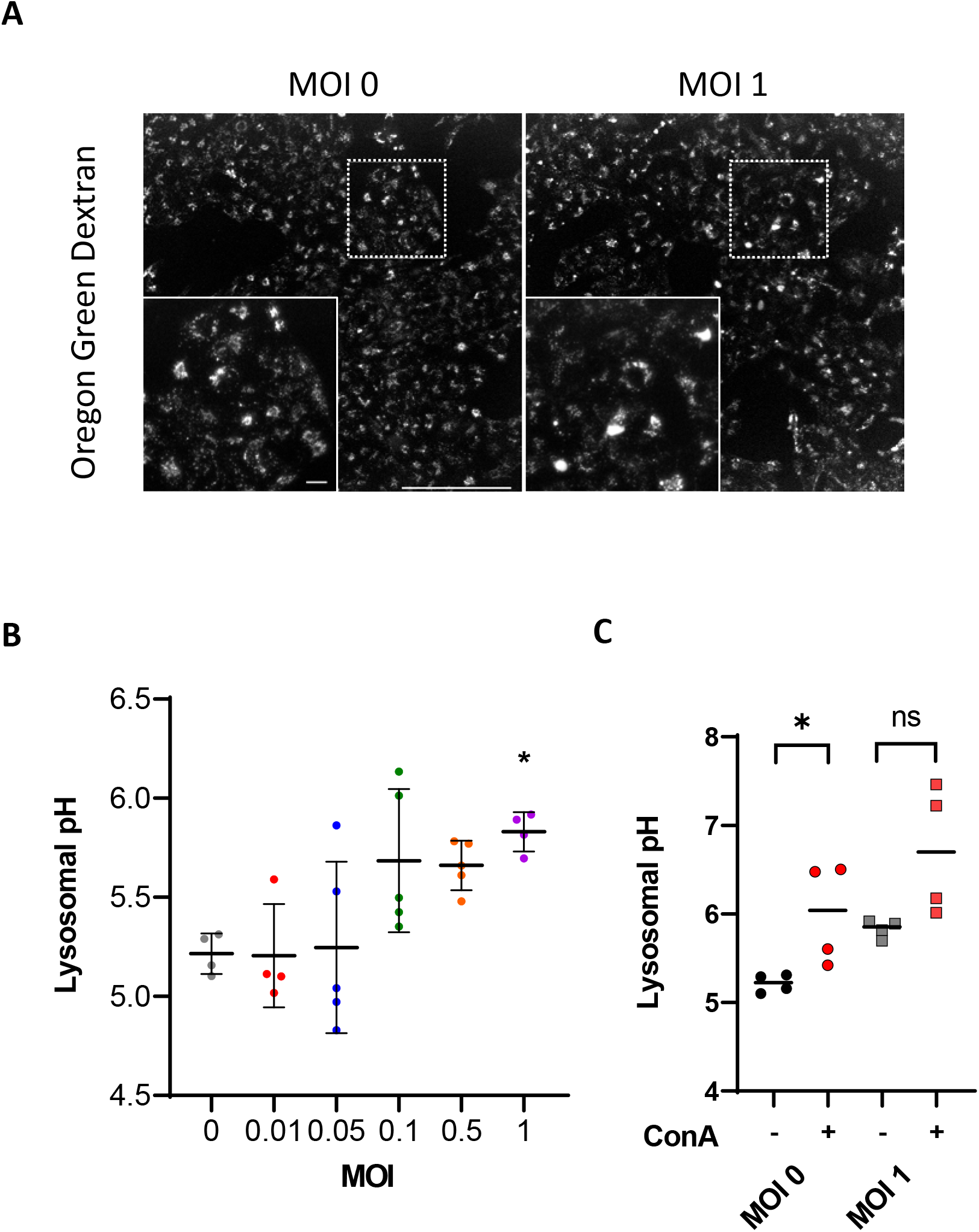
Effect of SARS-CoV-2 infection on the lysosomal pH of Vero E6 cells. (A) Representative fluorescence images (λex=488, λem=510) of VeroE6 cells loaded overnight with OGDx and infected with 0 or 1 MOIs of the Delta SARS-CoV2 virus. Bar size 200 μm and 20 μm (inset). (B) Lysosomal pH of Vero E6 cells infected with different MOIs of the Delta SARS-CoV2 virus. Each dot shows the averaged pH of 40-60 cells from 4-5 fields in one of two independent experiments, lines show mean±SD. *p<0.05, ns, non-significant, ordinary one-way ANOVA. (C) Lysosomal pH of Vero E6 cells infected with 0 and 1 MOIs of the Delta SARS-CoV2 virus, treated or not with ConA (1μM, 10min). Each dot shows the averaged pH of 15-50 cells from 4 wells in one of two independent experiments, lines show median values. *p<0.05, ns, non-significant, two tailed unpaired Student T test.

### The envelope protein of SARS-CoV-2 accumulates in the ERGIC when expressed in Vero E6 cells

To test whether the pH dissipating effects of the SARS-CoV-2 virus reflect the activity of its E protein in organelles, we transiently expressed plasmids coding for the native and epitope-tagged E protein in mammalian Vero E6 cells. Expression of wild-type SARS-CoV-2 E in Vero E6 cells was confirmed by the detection of a ~15 kD band on western blots with an in-house generated recombinant antibody directed against the native protein (Fig. 4A). A band of similar size was detected in cells expressing E mutated at residues N15 and V25 within the putative pore domain, and with antibodies against the streptavidin epitope (WSHPQFEK) added to the C terminus of the protein (Fig. 4A), confirming expression of the recombinant proteins. Streptavidin immunoreactivity was detected in Vero E6 cells expressing Strep-tagged E in structures overlapping with ERGIC-53 immunoreactivity and with co-expressed sec22b-rpHl (Fig. 4B). When expressed in HeLa cells, Strep-tagged E colocalized extensively with GFP-ERGIC-53 and partially with the ER marker RFP-KDEL (Fig. S4A), and the pattern of these ER and ERGIC markers was not altered by the co-expression of the E protein (Fig. S4B). These data indicate that the ectopically expressed viral E protein accumulates preferentially in the ERGIC.

**Figure 4.**
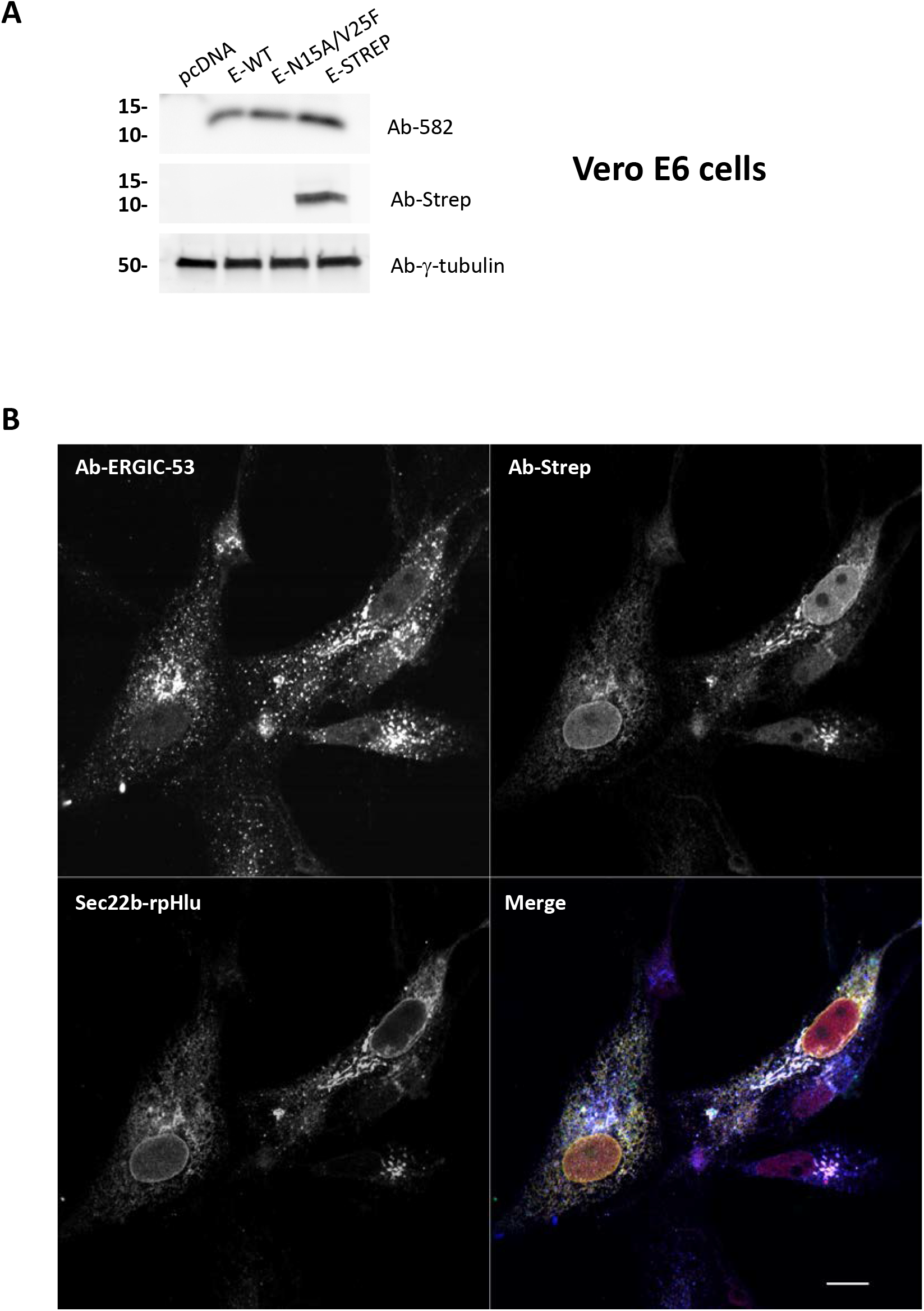
Expression of the envelope protein of SARS-CoV-2 in Vero E6 cells. (A) Western blots of whole-cell lysates from Vero E6 cells transiently expressing the empty vector (pcDNA), the native SARS-CoV-2 E protein (E-WT), the N15A/V25F SARS-CoV-2 E double mutant (E-N15A/V25F), and the streptavidin-tagged SARS-CoV-2 E protein (E-Strep). Blots were revealed with a recombinant antibody raised against the native SARS-CoV-2 E protein (Ab-582, top), an anti-streptavidin antibody (middle) or with γ-tubulin as a loading control (bottom). (B) Fluorescence images from VeroE6 cells transiently expressing Sec22b-rpHluorin (green) together with Strep-tagged SARS-CoV2 E protein (red, streptavidin immunoreactivity), stained with an anti-ERGIC53 antibody (blue). Bar size 10μm.

### The envelope protein of SARS-CoV-2, but not its pore mutant, dissipates the ERGIC pH

We then measured the impact of E protein expression on the ERGIC and cytosolic pH. Acute expression of the wild-type E protein (E-WT) increased the resting pH_ERGIC_ of Vero E6 cells from 7.03±0.04 to 7.19±0.03 (Fig. 5A). In contrast, expression of a E protein mutated at residues N15 and V25 within the pore domain did not impact the ERGIC pH (Fig. 5A). In all conditions, the pH of the ERGIC was significantly lower than the resting cytoplasmic pH of 7.29±0.10 (Fig. 5A). To assess the cellular toxicity associated with the expression of E, we measured cell death and ER stress in HEK-293 cells. Short-term (24h) expression of E bearing or not the V25F channel mutation did not induce apoptosis or ER stress, while longer expression (72h) increased CHOP mRNA levels and XBP1 splicing, the latter being only observed in cells expressing the WT protein (Fig. S5). These data indicate that the presence of the envelope protein in the ERGIC dissipates the acidic pH of this intermediate compartment. This viroporin effect is associated with delayed ER stress responses but not with acute cellular toxicity and is prevented by mutations within the channel pore of E.

**Figure 5.**
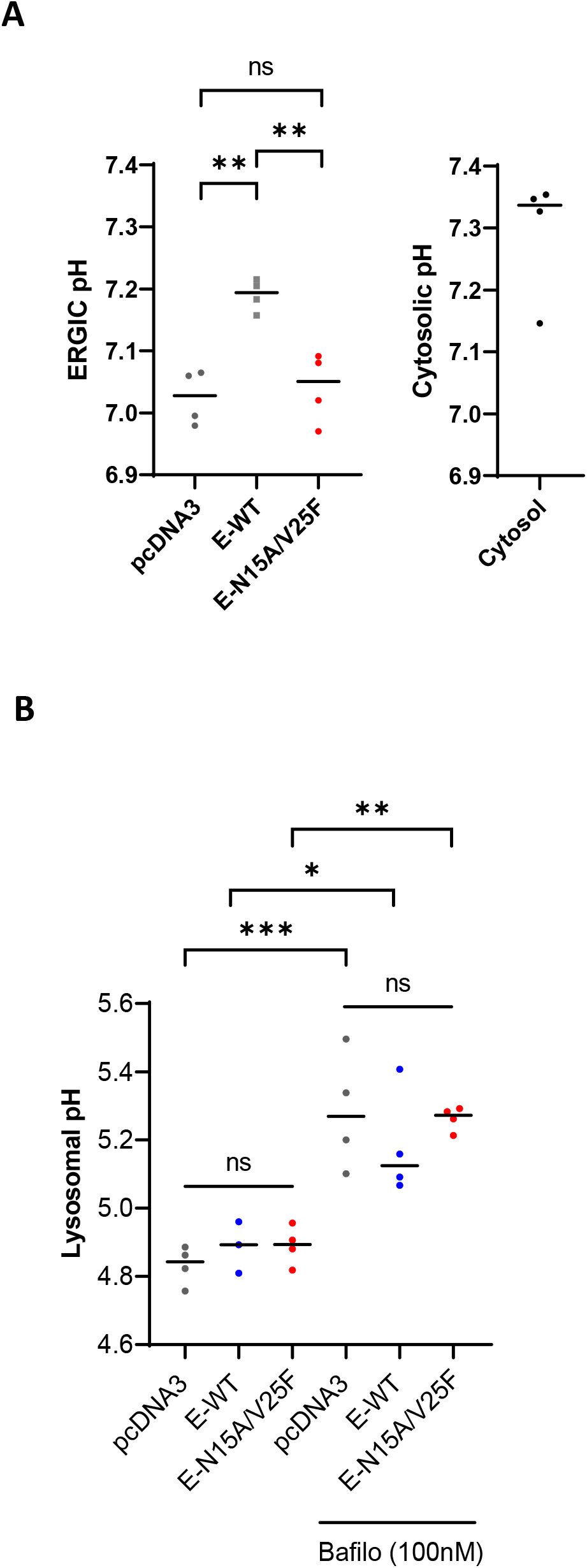
Effect of SARS-CoV-2 E on the cytosolic and organelle pH. (A) ERGIC pH (left) and cytosolic pH (right) of Vero E6 cells transiently expressing the empty vector (pcDNA3), the native SARS-CoV2 E protein (E-WT) and the N15A/V25F SARS-CoV2 double mutant (E-N15A/V25F). Each dot shows the average pH of 30-80 cells from an experiment, lines show median values. Ns, non-significant, **p<0.01. Ordinary one-way ANOVA. (B) Lysosomal pH of Vero E6 cells transiently expressing the indicated constructs, treated or not with Bafilomycin A1 (100 nM, 10 min). Each dot shows the average of 30-80 cells from one of 4 replicates in 2 experiments, lines show median values. Ns, non-significant, *p<0.05, **p<0.01,***p<0.005. Ordinary one-way ANOVA.

### The envelope protein of SARS-CoV-2 does not affect the resting pH of lysosomes

We then measured the lysosomal pH (pHlyso) of Vero E6 cells with OGDx. With the widefield microscope we were able to measure the dextran fluorescence ratio (λex=430/490, λem=530) and generated a titration curve for the conversion of experimental ratio to pH (Fig. S3A). pHlyso was not altered by the expression of the wild-type or mutated envelope protein (Fig. 5B). Addition of the V-ATPase inhibitor bafilomycin alkalinized lysosomes by ~0.2 pH units regardless of E protein expression (Fig. 5B). These data indicate that the expression of the envelope protein of SARS-CoV-2 specifically dissipates pH in ERGIC compartment but does not affect the acidic pH of lysosomes.

## Discussion

In this study, we report that infection of Vero E6 cells with the SARS-COV-2 delta virus alkalinizes the ERGIC and lysosomes and link this pH alteration to the ion channel function of the viral envelope protein in the ERGIC. Using the ratiometric pHluorin fused to Sec22b, we provide the first direct quantitative measurements of the pH within the lumen of the ERGIC. We found that the ERGIC pH of Vero E6 cells is ~0.2 pH units more acidic than the cytosolic pH, measured with the same genetic indicator, due to the activity of vacuolar H^+^-ATPases sensitive to bafilomycin and concanamycin. SARS-COV-2 infection, at MOI higher than 0.1, dissipated the acidic pH of the ERGIC and lysosomes, mimicking the effects of V-ATPases inhibitors. In non-infected cells, enforced expression of the SARS-COV-2 E protein dissipated ERGIC pH but did not alter lysosomal pH, while expression of a channel pore mutant did not impact ERGIC pH. These data establish that E acts as a proton channel within the ERGIC and that the SARS-COV-2 virus exploits this viroporin activity to alter the pH of intracellular organelles.

Recent electrophysiological recordings of HEK-293 cells and Xenopus oocytes expressing SARS-COV-2 E targeted to the plasma membrane reported large monovalent cations currents that became inward rectifying as the pH was decreased from 8 to 6 in HEK-293 cells (Cabrera-Garcia et al., 2021). This indicates that the E protein forms a cation channel activated at luminal acidic pH, as previously reported for other coronaviruses (Mandala et al., 2020). These authors further showed that SARS-COV-2 E fused to mKate accumulates in perinuclear structures decorated by an anti-ERGIC-53 antibody and decreases the fluorescence of a membrane-permeant pH-sensitive dye (Lysosensor DND-189, pKa=5,2) in NIH-3T3 cells (Cabrera-Garcia et al., 2021). This indicates that enforced E expression increases the luminal pH of acidic organelles, and effect previously reported for the viroporins of other coronaviruses (Westerbeck and Machamer, 2019)

We confirm here that the SARS-COV-2 E protein accumulates in the ERGIC when ectopically expressed in mammalian cells. In Vero E6 cells, the E protein colocalized extensively with the mannose-specific membrane lectin ERGIC-53, an established ERGIC marker, and with rpHluorin fused to the transmembrane domain of the ERGIC-resident SNARE Sec22b. In HeLa cells, the epitope-tagged SARS-CoV-2 E protein was detected predominantly in the ERGIC with a minor fraction in the ER. Acute E expression did not impact membrane integrity or organelle appearance and did not induce ER stress at 24 or 48 hours, but only at 72 hours. Enforced E expression, on the other hand, increased the luminal pH of the ERGIC by 0.2 pH units in Vero E6 cells without altering the pH of lysosomes, measured by internalized OGDx. ERGIC alkalinization was not observed in cells expressing an E mutant bearing two amino acids substitutions within the predicted pore domain, linking the alkalinization to the proton channel function of E. Importantly, a similar viroporin activity was detected in cells infected with a replicating SARS-COV-2 delta virus. Viral infection alkalinized the ERGIC and lysosomes, mimicking the effects of V-ATPases inhibitors. The simplest explanation for the ERGIC alkalinization occurring in infected cells is therefore that the viral envelope protein acts as a viroporin in this organelle. The lysosomal alkalinization observed in infected cells might also reflect the viroporin activity of E, which could reach this compartment in the full virus.

Our report that SARS-CoV-2 infection alter the ERGIC and lysosomal pH of mammalian cells has implications for virus fitness and pathogenicity. The SARS-COV-1 E protein accumulates in the ER and Golgi and its viroporin activity in these organelles is thought to facilitate virus propagation and pathogenicity (DeDiego et al.; Nieto-Torres et al.). We show that the proton channel function of SARS-COV-2 E counteracts ERGIC acidification by vacuolar ATPases. Mitigating ERGIC and lysosome acidification might protect newly formed virions from a toxic acidic environment. An acidic pH is important for dissociation of cargo from sorting lectins such as ERGIC-53 (Appenzeller-Herzog et al., 2004) and SARS-CoV-2 might exploit this mechanism to promote viral particle assembly by relying on the viroporin activity of E to dissipate the pH gradient. Preventing organelle acidification is also expected to disrupt secretory cargo relying on pH-sensitive dissociation mechanism, like procathepsin. The ion channel function of the SARS-CoV-2 E protein therefore likely contributes to SARS-CoV-2 propagation and pathogenicity, making this viroporin an attractive drug target.

In summary, we show here that SARS-CoV-2 infection prevents ERGIC and lysosomes acidification in infected cells and link the ERGIC pH deregulation to the viroporin activity of the viral envelope protein. The channel function of E likely contributes to SARS-CoV-2 fitness and pathogenicity by alkalinizing organelles. Compounds inhibiting this viroporin could provide new antiviral drugs targeting an essential viral function conserved among coronaviruses.

**Supplementary Figure 1.**
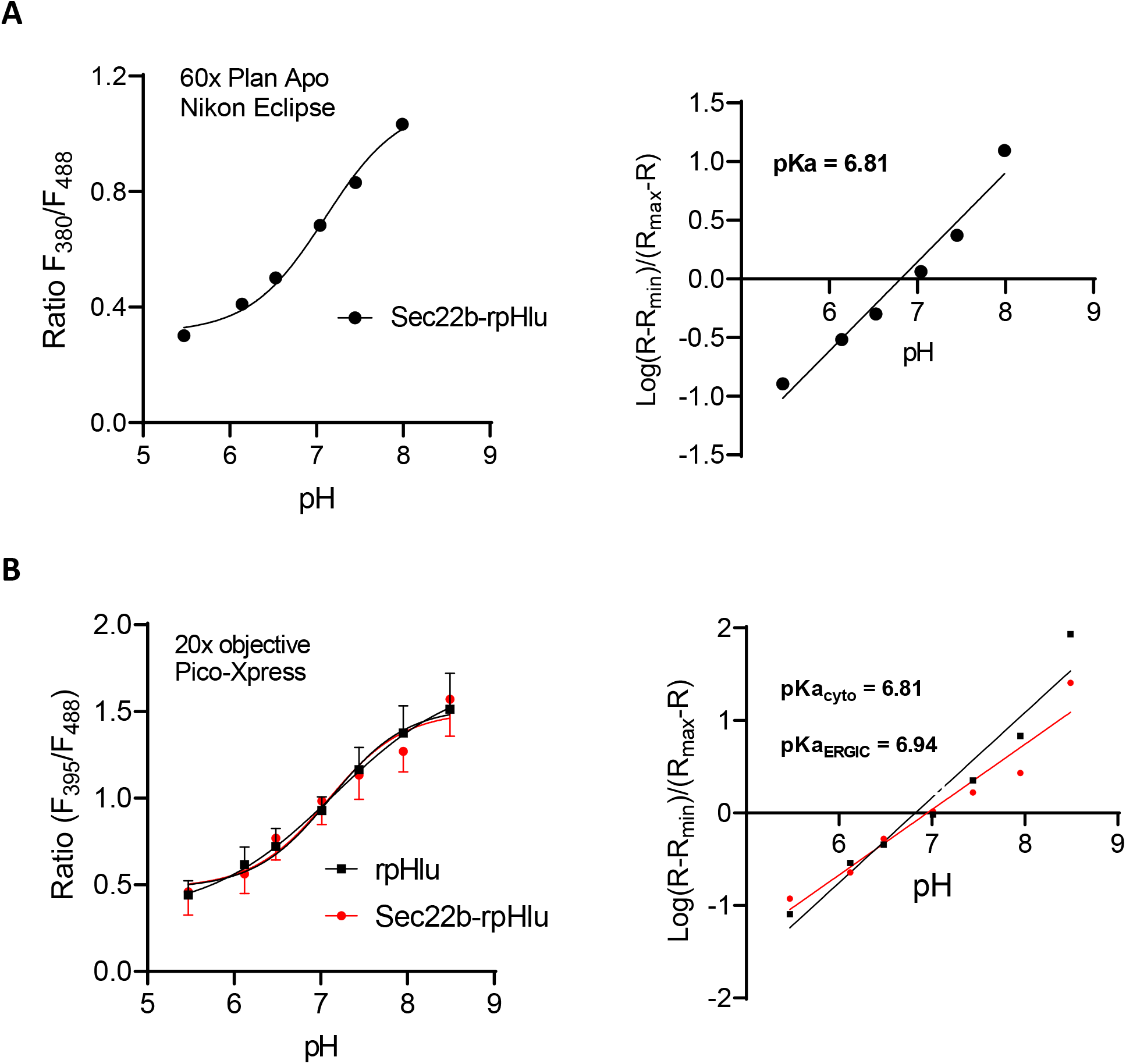
Calibration of cytosolic and ERGIC-targeted rpHluorin. (A) Left: *In-situ* pH titration of sec22b-rpHluorin fluorescence ratio on a high-resolution fluorescence microscope (λex=380/488, λem=510). Each dot is the average of 40-50 cells from 2 experiments, each with 15-20 image fields. Line is a sigmoidal fit of the data. Right: log-log display of the pH titration curve. Line is a linear fit of the data, crossing the x axis at the probe’s pKa. (B) *In-situ* pH titration of rpHluorin and sec22b-rpHluorin fluorescence ratio on an automated cell imaging system (λex=395/488, λem=510). Each dot is the average of 40-50 cells from 2 experiments, each with 15-20 image fields. Lines are sigmoidal fits of the data. Right: log-log display of the pH titration curves with linear fits that cross the x axis at nearly identical values corresponding to the probe’s pKa.

**Supplementary Figure 2.**
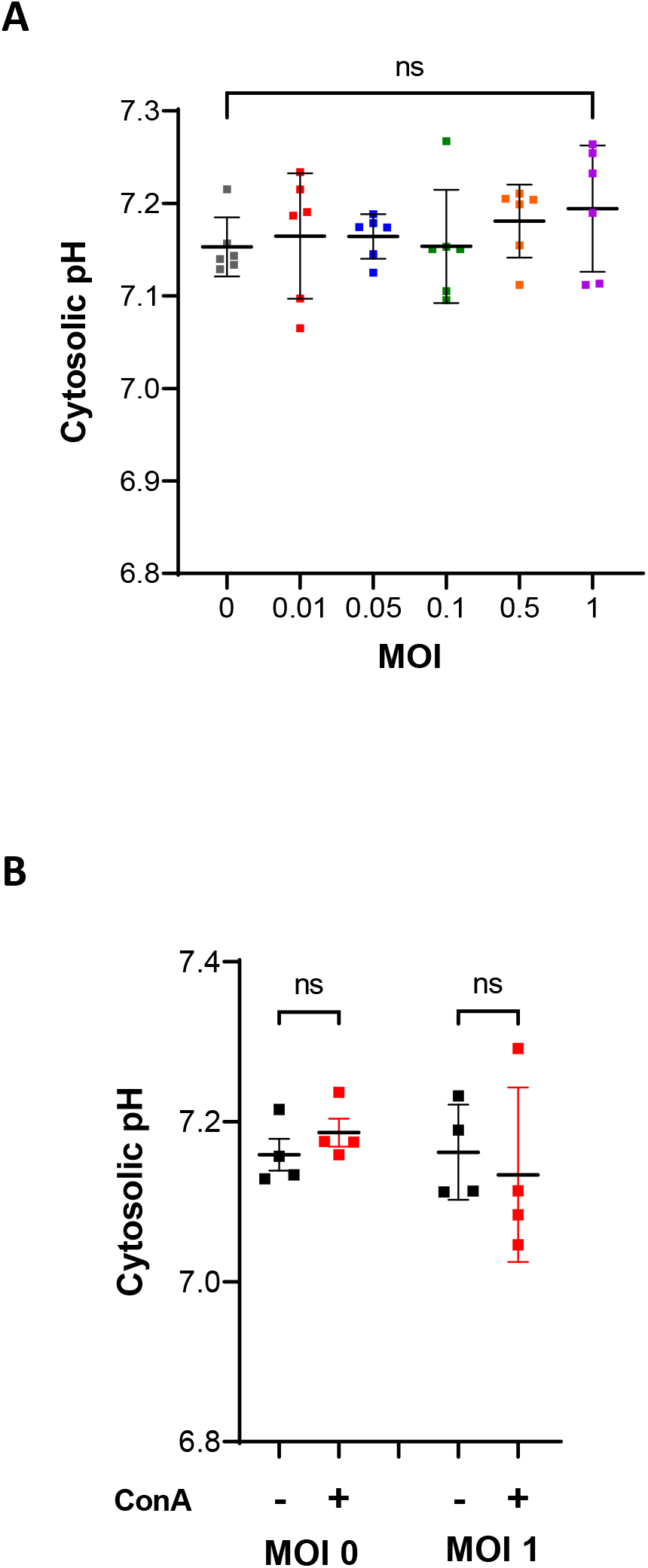
Effect of SARS-CoV-2 infection on the cytosolic pH of Vero E6 cells. (A) Cytosolic pH of Vero E6 cells infected with different MOIs of the Delta SARS-CoV2 virus. Each dot is the average of 10-30 cells from 3 independent experiments performed in duplicates, lines are mean±SD. *p<0.05, ns, non-significant, ordinary one-way ANOVA. (B) Cytosolic pH of Vero E6 cells infected with 0 (data from Fig. 1C) and 1 MOIs of the Delta SARS-CoV2 virus and treated or not with ConA (1μM, 10min). Each dot shows the averaged pH of 15-50 cells from 4 replicates in one of two independent experiments, lines mean±SD. *p<0.05, ns, non-significant, ordinary one-way ANOVA.

**Supplementary Figure 3.**
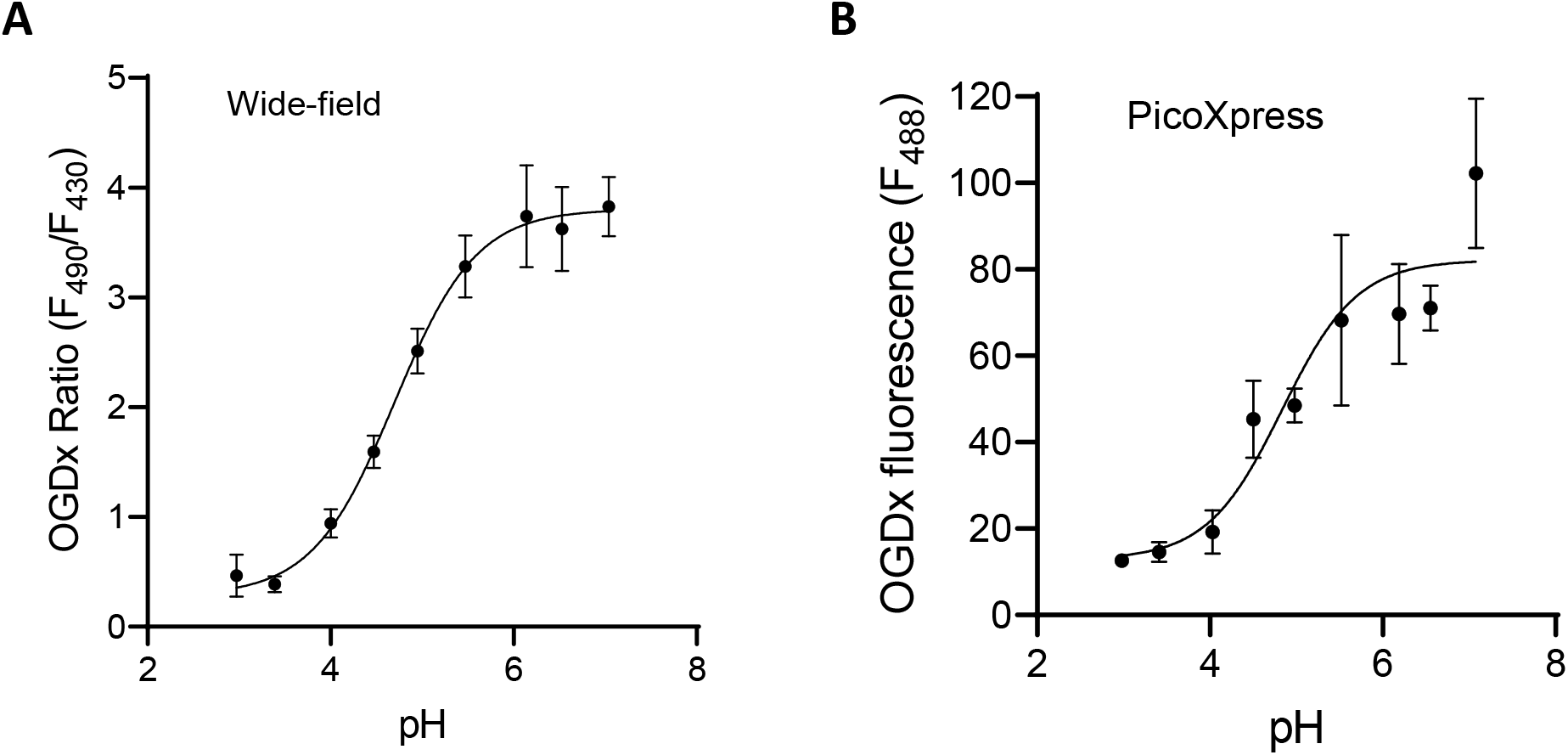
Calibration of OGDx internalized in lysosomes. (A, B) *In-situ* pH titration of internalized OGDx-488 ratio fluorescence (λex=430/490, λem=530) on a high-resolution microscope (A) and of OGDx-488 single fluorescence (λex=490, λem=530) on the automated cell imaging system (B). Each dot shows the average fluorescence of 20 cells from 4-5 fields in one of 2 independent measurements. Lines show sigmoidal fits of the data.

**Supplementary Figure 4.**
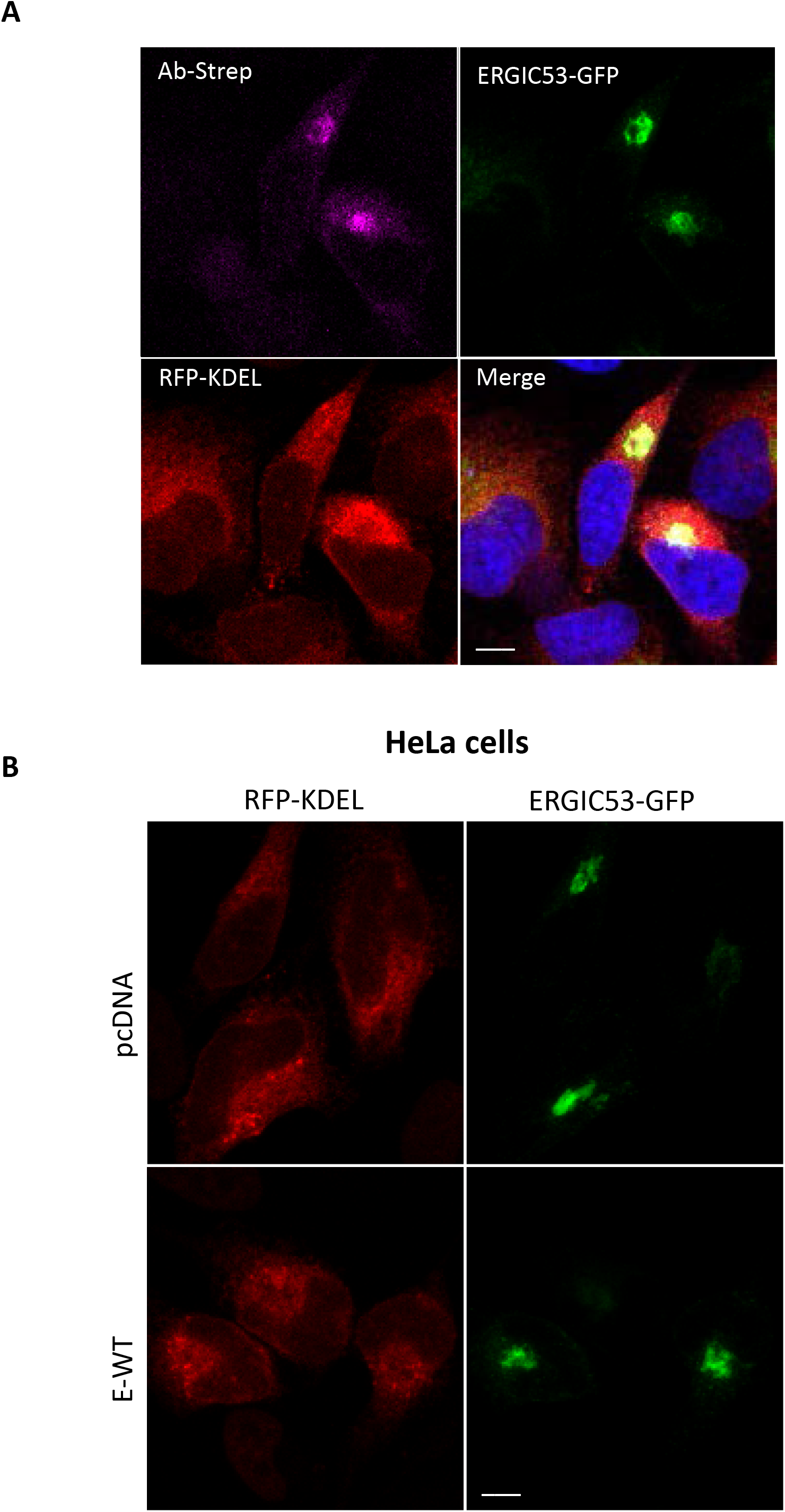
Expression of SARS-CoV-2 E in HeLa cells. (A, B) Fluorescence images of HeLa cells transiently expressing Strep-tagged E-WT (violet) together with ERGIC53-GFP (green) and RFP-KDEL (red). Bar size 10μm.

**Supplementary Figure 5.**
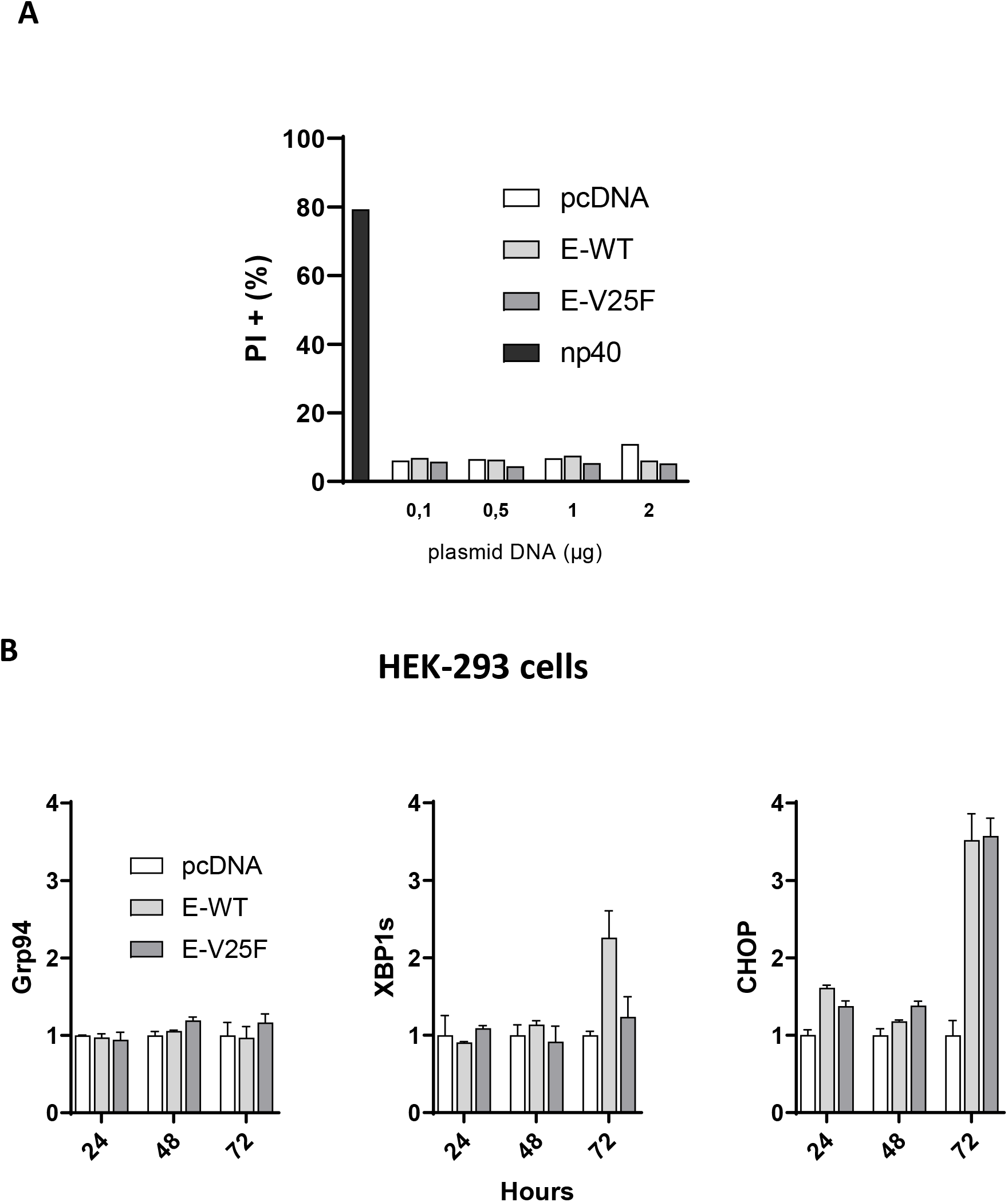
Effects of SARS-CoV-2 E expression in HEK-293 cells. (A) Percentage of propidium iodide (PI) positive cells determined by flow cytometry in HEK-293 cells transiently expressing pcDNA3, E-WT, and E-V25F for 24h. (B) qRT-PCR analysis of mRNA transcripts of CHOP, Grp94, and spliced XBP1, normalized to the expression of HPRT mRNA in HEK-293 cells expressing pcDNA3, E-WT, and E-V25 for up to 3 days. Data are mean ±SEM of two independent experiments in triplicate.

## Materials and Methods

### Reagents

Concanamycin A was purchased from Enzo-Life-Sciences (AXL-380-034-C100), Bafilomycin A1 from Sigma (B1793). The mouse anti SARS-CoV-2 E protein nanobodies were developed by the Geneva antibody facility and described in DOI: https://doi.org/10.24450/journals/abrep.2020.e191, the mouse anti STREP-Tag antibody was purchased from BioLegend (688202), the mouse anti gamma-tubulin antibody from ThermoFisher (MA1-850) and the rabbit anti ERGIC-53 antibody from MERCK (E1031). The SARS-CoV-2 E-protein wild-type and N15A/V25F cDNA sequences were ordered as synthetic plasmids with flanking EcoRI and XhoI cut sites from GeneArt and cloned into pcDNA using EcoRI and XhoI enzymatic digestion. The pTwist-EF1a carrying the SARS-CoV-2 E protein with 2xSTREP tags were obtained from Addgene. pCMV-rpHluorin-N1 was a kind gift from Dr. Thierry Galli (INSERM, Paris). The pCMV-sec22b-epHluorin was generated by cloning sec22b into the multiple cloning sites of pCMV-rpHluorin-N1, using Nhei and XhoI enzymatic digestion.

### Cells and transfection

Vero-E6 cells were a kind gift from Pre Caroline Tapparel (UNIGE, Geneva) and Calu-3 cells were a kind gift from Dr. Karl-Heinz Krause (UNIGE, Geneva). Vero E6 cells were cultured in Dulbecco’s Minimal Essential Medium (DMEM) supplemented with 10% FBS and Pen/Strep and maintained at 37°C and 5% CO2. Cells were grown to 80% confluency prior to transfection or co-transfection with different plasmids for 24 hours using lipofectamine 2000 (ThermoFisher).

### Viral preparation and infection

The delta-SARS-CoV2 virus was a kind gift from Dr. Isabella Eckerle (University Hospital Geneva). For propagation, the virus was cultured with Calu-3 cells for 72 hours prior to collection of media and clarification of cell debris by centrifugation (2000 rcf). The pfu of the virus was 10e6, determined by Vero E6 cell infection followed by plaque assay in a 24 well plate format, using cells plated to 80-90% in a 24 well plate and were infected with serial dilutions of the virus.

### pH measurements

For cytoplasmic and ERGIC pH recordings, Vero E6 cells were transfected with pCMV-rpHluorin-N1 or pCMV-sec22b-rpHluorin. Cells transiently expressing or not the envelope protein were alternatively excited for 100 ms with ET380x and ET490/20 filters and rpHluorin ratio fluorescence imaged with a 525/50 band pass filter (Chroma) at 37°C on a Nikon Eclipse Ti inverted microscope equipped with a 60x Plan Apo 1.30NA objective; Sutter Lambda XL lamp; Bipolar Temperature control stage heater (Harvard Apparatus); controlled by Visiview software (Visitron Systems). Cells infected with the virus were imaged inside a BSL3 lab on a PicoXpress microscope (Molecular Devices) on 96 well plates alternatively excited for 500ms through the FITC 445-485/509-539 filter cube and 800ms through the customized F49-395 395/25ET Bandpass; F48-425 Beamsplitter T 425 LPXR; F47-525 525/50 ET Bandpass filter cube on the HC PL FLUOTAR 20x/0.40 objective.

For lysosomal pH, Vero E6 cells were loaded overnight with Oregon Green™ 488, dextran (OGDx) 10,000MW (D7171, ThermoFisher) and prepared and imaged as previously described (Pihan et al., 2021). Cells infected with the virus were imaged with the PicoXpress in the BSL3 lab, using a single excitation for 1000ms through the FITC 445-485/509-539 filter cube.

pH calibration was performed using nigericin (5 μg/ml) and monensin (5 μM) in solutions containing 125 mM KCl, 20 mM NaCl, 0.5 mM MgCl2, 0.2 mM EGTA, and HEPES (pH 7.0-7.5), or MES (pH 5.5–6.5), or acetic acid (pH 4-4.5) or citric acid (pH 3-3.5) as previously described (Nunes et al., 2015; Pihan et al., 2021). The cells were incubated with each calibration solution for 3 minutes before imaging. For each experiment, a five-point calibration curve was fitted to a variable slope sigmoid equation using GraphPad Prism. Cells were imaged in modified Ringer’s buffer or in Vero E6 culture media supplemented with 25 mM HEPES.

### Cell death and qPCRs

HEK cells were transfected with the indicated plasmids and plated for the indicated times before imaging propidium diiodide accumulation in a Fortessa II flow cytometry machine. For ER stress, RNA was harvested at the indicated times and qPCRs were performed with the following primers (5’-3’): GRP94 TCCATATTCGTCAAACAGACCAC; CTGGGACTGGGAACTTATGAATG; XBP1s TGCTGAGTCCGCAGCAGGTG; GCTGGCAGGCTCTGGGGAAG; CHOP ATTGACCGAATGGTGAATCTGC; AGCTGAGACCTTTCCTTTTGTCTA, HPRT TGACACTGGCAAAACAATGCA; GGTCCTTTTCACCAGCAAGCT (Forward and reverse respectively). Data was normalized to HPRT

### Western blotting

Following transfection, cells were harvested with RIPA lysis buffer (Sigma; R0278) containing protease inhibitor (Sigamafast^TM^ protease inhibitor cocktail tablets, EDTA-free) for 30 minutes on ice. Cell lysates were centrifuged at 11,200 x g for 10 minutes and the supernatant was diluted with 4X NuPAGE LDS Sample Buffer (ThermoFisher; NP0007). Samples were subjected to electrophoresis through 4-20% mini-protean® TGX™ precast gels (BioRad; 4561095), membrane transfer and immunoblot analysis. Immunoblots were probed with mouse anti SARS-CoV-2 E protein nanobody (1:250), mouse anti STREP-Tag (1:1000) and mouse anti γ-tubulin (1:2000).

### Immunofluorescence

Immunofluorescence was performed in Vero E6 cells co-transfected with the indicated constructs. After 24h of transfection, cells were fixed (Pfa 4%) for 20 min at room temperature, then permeabilized (0.5% BSA in PBS + 0.5% Triton X-100) for 10 min at RT and blocked (2% BSA in PBS) for 1 hour at room temperature. Cells were then incubated with primary antibodies overnight at 4ºC in a wet chamber and then incubated with the corresponding secondary antibodies coupled to fluorochromes (1:1000) for 1h at RT. Images were obtained on a LSM700 Nikon microscope.

### Image analysis and statistics

Image analysis was performed with ImageJ. Data analysis was performed with GraphPad Prism 8. Student’s T-test and ANOVA statistical analysis were used where appropriate.

## Data availability

The data that support the findings of this study are available from the corresponding author upon reasonable request.

## Acknowledgements

We are grateful to the bioimaging core facility of the Faculty of Medicine of the University of Geneva. The graphical abstract was created with BioRender.com. This work was funded by the Swiss National Foundation [grant number 310030_189042 (to N.D.)

## Author contributions

WAW, Acquisition of data, Analysis and interpretation of data, Drafting or revising the article; ND, Conception and design, Analysis and interpretation of data, writing of the article.

